# Glucose dynamics during ozone exposure measured using radiotelemetry: Stress drivers

**DOI:** 10.1101/2021.12.09.471963

**Authors:** Andres R. Henriquez, Samantha J. Snow, Thomas W. Jackson, John S. House, Alison A. Motsinger-Reif, Cavin K. Ward-Caviness, Mette C. Schladweiler, Devin I. Alewel, Colette N. Miller, Aimen K. Farraj, Mehdi S Hazari, Rachel Grindstaff, David Diaz-Sanchez, Andrew J Ghio, Urmila P. Kodavanti

## Abstract

**Background:** Stress-related neurobehavioral and metabolic disorders are associated with altered circulating adrenal-derived hormones and hyperglycemia. Temporal assessment of glucose and these hormones is critical for insights on an individual’s health.

**Objectives:** Here we used implantable-telemetry in rats to assess real-time changes in circulating glucose during and after exposure to the air pollutant ozone, and link responses to circulating neuroendocrine stress and metabolic hormones. We also compared rodent glucose and corticosterone (cortisol in humans) responses to humans exposed to ozone.

**Methods:** First, using a cross-over design, we monitored glucose levels during single or repeated ozone exposures (0.0, 0.2, 0.4 and 0.8-ppm) and non-exposure periods in male Wistar-Kyoto-rats implanted with glucose-telemeters. A second cohort of un-implanted rats was exposed to ozone (0.0, 0.4 or 0.8-ppm) for 30-min, 1-hour, 2-hour, or 4-hour with hormones measured immediately after exposure. Then we assessed glucose metabolism in sham and adrenalectomized rats with or without pharmacological interventions of adrenergic and glucocorticoid receptors. Finally, we assessed glucose and cortisol in serum samples from a clinical study involving exposure of human volunteers to air or 0.3 ppm ozone.

**Results:** Ozone (0.8-ppm) caused hyperglycemia and hypothermia beginning 90-min into exposure, with reversal of effects 4-6 hours post-exposure. Glucose monitoring during four daily 4-hour ozone exposures revealed duration of hyperglycemia, adaptation, and diurnal variations. Ozone-induced hyperglycemia was preceded by increased adrenocorticotropic hormone, corticosterone, and epinephrine, but depletion of thyroid-stimulating, prolactin, and luteinizing hormones. Hyperglycemia was inhibited in rats that were adrenalectomized and/or treated with glucocorticoid inhibitor. The depletion of cortisol was dampened in humans exposed to ozone during intermittent exercise.

**Discussion:** We demonstrate for the first time the temporality of neuroendocrine-stress-mediated biological sequalae responsible for ozone-induced metabolic dysfunction as exposure occurs. Real-time glucose monitoring with stress hormones assessment may be useful in identifying interactions among pollutants and stress-related illnesses.

## INTRODUCTION

Air pollution, climate change, the epidemic of COVID-19, social inequalities, unhealthy dietary habits, and sedentary lifestyle are likely to further escalate mental health crises and metabolic syndrome world-wide. Among many environmental risk factors, air pollution accounts for nearly 70% of all environmental causes of human mortality^1^, and is linked to neurobehavioral and metabolic diseases. Increased incidence of Alzheimer’s disease,^2,3^ late life cognitive decline,^4^ anxiety, and even criminality have been associated with the exposure to air pollutants.^5^ Moreover, associations have been found between air pollution and concurrent exacerbation of diabetes and Alzheimer’s disease.^4^ Those with Alzheimer’s and Parkinson’s disease also often suffer from diabetes suggesting potential neural contribution to peripheral diseases and mechanistic linkages.^6,7^

With the emerging link between air pollution, stress, and neuro-cognition,^5,8^ the role of neuroendocrine system deserves further exploration. Psychosocial and environmental stressors are the primary contributors to chronic disease susceptibility. The central neuroendocrine system responds to stress-induced autonomic sensory activation and orchestrates a peripheral stress response, while initiating two survival processes, namely metabolism and immune surveillance in an organ-specific manner.^9^ In addition to sympathetic neurons innervating the majority of peripheral organs, sympathetic-adrenal-medullary (SAM) and hypothalamic-pituitary-adrenal (HPA) axes-mediated release of catecholamines and corticosteroids produce peripheral cellular response to stress.^9^ These are the same hormones that regulate stress adaptation through their action on the central nervous system (CNS) centers via feedback controls.^10^ Chronic alterations in the levels of circulating stress hormones, especially glucocorticoids, have been linked to psychological disorders and cardiometabolic diseases.^11^ Centrally-mediated release of stress hormones and peripheral responses to stress are plastic, and are temporally and spatially regulated such that no adverse effects persist upon discontinuation of stress.^10^ However, when the neuroendocrine system is impaired or overactive, disease may ensue.^12^ Thus, the temporal assessment of the stress dynamics is critical to understanding an individual’s susceptibility to environmental insults.

Monitoring of stress response through real-time assessment of circulating cortisol and glucose in humans is not common. However, in children recovering from surgery, blood glucose and cortisol are often measured to assess stress.^13^ Individuals with type 2 diabetes, when subjected to acute moderate psychological stress (Trier Social Stress Test), have spikes in blood glucose as determined using real-time glucose monitors.^14^ Real-time glucose monitoring sensors, which have been developed in early 1990’s, are now applied more widely for diabetics.^15^ In humans, continuous monitoring of glucose using non-invasive sensor-based electromagnetic coupling is already gaining popularity,^16^ providing the opportunity to use such monitors in detecting environmental stressor effects such as air pollutants. Very recently, a wireless graphene-based sweat stress sensing mHealth system has been developed for dynamic and non-invasive assessment of cortisol in sweat.^17^ The use of new techniques for dynamic stress assessment will be valuable for determining the health impact of stressors on the body. In particular, this approach in air pollution studies could allow evaluation of short and long-term effects on health and provide diagnostic and mechanistic insights.

We have shown that acute single exposure to ozone induces a classical stress response associated with increases in circulating catecholamines, glucocorticoids, and glucose.^18,19^ However, the neuroendocrine stress response is temporal and reversible in healthy individuals. To link ozone-induced alterations in metabolic processes and neuroendocrine stress, it is critical to determine the dynamicity of peripheral metabolic effects and how that relates to neuroendocrine changes. The purpose of our study was to determine the temporality of glucose changes during and after ozone exposure in a rat model using implantable radiotelemetry, which has not been previously employed for experimental assessment of air pollution effects. We hypothesized that increased glucose levels during ozone exposure will be mediated by ozone-induced increases in stress hormones. Real-time glucose measures were coupled with separate measures of stress hormones as well as mechanistic studies to assess the role of stress responses in ozone-induced glucose fluctuations. Moreover, to establish coherence with human responses, we assessed the impacts of ozone exposure on glucose and cortisol levels in young healthy volunteers.

## Materials and Methods

### Animals

All male Wistar Kyoto rats were purchased from Charles River Laboratories Inc., Raleigh, NC at 10-12 weeks of age and maintained in our Association for Assessment and Accreditation of Laboratory Animal Care-approved EPA animal facility. Animals were pair-housed in polycarbonate cages with beta chips bedding and EnviroDry enrichment material, except when stated. Animal rooms were maintained on a 12 hr light/dark cycle (6AM-6PM) at ∼22°C and 50% relative humidity. They were provided free access to Purina (5001) pellet rat chow (Brentwood, MO) and water ad libitum, unless stated during experimental procedures. All animal protocols were approved by the EPA’s Institutional Animal Care and Use Committee prior to starting studies and we followed National Institutes of Health guide for the care and use of rats (NIH Publications No. 8023).

### Glucose telemetry surgeries

Eight male Wistar-Kyoto (WKY) rats (at 13 weeks of age) were implanted with DSI glucose telemeters (HD-XG, St. Paul, MN) using aseptic techniques (Figure S1, Table S1). Anesthesia was induced by vaporized isoflurane inhalation (4%, 1-2 LPM of O_2_) and maintained during the surgery (2-3%, 1-2 LPM of O_2_). Once anesthetized, analgesic meloxicam (2 mg/kg, in saline s.c.) and artificial tear ointment were provided before surgery. Anesthesia was continuously checked by toe pinch. A trained surgeon implanted the sensor in the descending abdominal aorta and the transmitter subcutaneously. The blood glucose (Nova Biomedical, Waltham, MA) sensor uses glucose oxidase to convert glucose and oxygen into gluconic acid and hydrogen peroxide. The amount of hydrogen peroxide, which is proportional to the amount of glucose, reacts with a noble metal electrode to transfer electrons and create a current.^20^ The glucose telemetry system includes a reference electrode with an electronics/battery (Ag/AgCl) with a separate lead. This was sutured to the inner abdominal wall module into the midline abdominal muscle (Figure S1). During abdominal wall closure, the abdominal cavity was washed with saline (15 mL/kg) and rats were given bupivacaine (1 mg/kg, in saline s.c.). Recovery took place on heating pads under close observation of distress signals; once awake, the rats were placed into their home cages and administered with meloxicam (1 mg/kg, in saline s.c.) 24, 48 and 72 hours after the surgery.

### Blood glucose concentration data acquisition prior to exposure

After surgery, rats were individually housed in cages with pine shavings bedding. Rats were provided with water and powdered as well as pelleted standard Purina (5001) rat chow (Brentwood, MO) *ad libitum*. Blood glucose levels, core body temperature, and activity level measurements were sent via radio signals and collected using receivers placed under each cage. The recording for each rat began soon after the surgery and continued until the end of the experiment (∼ 9 weeks). The cages were placed over receivers (RPC-1, DSI) and data were simultaneously collected in a computer placed in an adjacent isolated room (Figure S1). For verification and calibration purposes, single point calibrations were carried out. As recommended, these calibrations were done twice weekly throughout the study based on the protocol explained by Brockway and collaborators at Data Sciences Inc.^20^ Data used for calibration are excluded from analysis and graphs. Interpolation of glucose levels and electric current detected by the sensor was corrected for differences in body temperature versus room temperature, merged and analyzed for each animal with a resolution of 1 minute using Dataquest® software acquisition system (DSI). Dataquest® acquisition system (DSI) included telemetry signal receiver for each animal, a data matrix for analyzing receiver signals and a computer. These telemetry devices allowed continuous sampling of 8 animals that are individually housed in cages with receivers directly underneath. To avoid signal mixing between two animals, the animal racks were equipped with stainless steel dividers. The software allowed device configuration and the protocol for continuous data collection as described previously.^20^

### Timeline of telemetered rat experiments

Two weeks after surgery, rats were exposed in pairs whole body to ozone (0.2, 0.8 or 0.4 ppm) or filtered air (0.0 ppm) in Rochester style “Hinners” chambers, using a crossover exposure design such that each rat was exposed to each concentration across the first 4 weeks of testing (Table S1). Briefly, rats for the first exposure were randomized in pairs for 0.0 (clean air), 0.2, 0.4 or 0.8 ppm of ozone exposure for 4 hrs. A different set of receivers were placed in each ozone exposure chamber to acquire data while animals were being exposed. After 1 week of washout period, these exposures were repeated but the targeted ozone exposure concentration for each pair of rats was changed. This experiment was repeated a total of four times over 4 weeks to cover all the targeted ozone concentrations for all eight rats (n=7 independent recordings). Rat #6 was discarded from the experiment due to malfunction of the sensor and/or transmitter. During week 5 and 6, these 7 telemetered rats were randomized into two groups, air and 0.8 ppm ozone, and again using crossover design they were exposed for 4 hours, once per week for 2 weeks (Table S1) and glucose levels were continuously monitored (n=7 per concentration). Immediately after each exposure, glucose tolerance test (GTT) was performed using a glucometer in addition to continuous recording of glucose through telemetry as described below. Finally, during week 7, animals were divided into air (n=3) and ozone (n=4) and at this time they were exposed to air or 0.8 ppm ozone 4 hours/day for four consecutive days to determine adaptation upon continuous exposure (Table S1). Because animals were exposed for 4 consecutive days during the week, a crossover design was not possible.

### Glucose/temperature telemetry data acquisition and analysis during exposures

During the day of exposure, animals were transported to the nearby exposure room where a control and three ozone exposure chambers were preprepared with a different set of receivers connected to an independent but identical data acquisition system to acquire data during exposure for each rat. Within ∼5 minutes of removal from their home cages, animals were immediately placed at designated locations (with receivers placed underneath) in the exposure chambers with wire-mesh bottom dividers and data were collected with the data acquisition system during exposure. Rats were place at two opposite corners of the chamber to avoid signal mixing. All exposures were aligned to minute 0 when 80% of target ozone concentration was reached. For each minute, glucose level and temperature were averaged from exposed rats (n=7/concentration). Immediately following the exposure, animals were likewise transported to their home cages for continued data acquisition. The data collected during non-exposure periods when telemetered rats are in their home cages and during air or ozone exposure periods were temporally aligned to tally continuous recording each day and during the entire experimental period. Glucose and temperature data with a resolution of 1 minute was averaged to prepare graphs. As mentioned above, for the adaptation study (7^th^ week), because of the daily exposure for 4 days, crossover design could not be used, and the sample size is thus n = 3 for 0.0 ppm and n = 4 for 0.8 ppm ozone. Multiple hours of data after the exposure stopped were included to follow days of recovery during non-exposure periods.

### Ozone generation and exposure for all rat experiments

Ozone exposure methods have been described in detail in previous publications.^18,21,22(p)^ Briefly, ozone was generated from oxygen using a silent arc discharge generator (OREC, Phoenix, Arizona) and measured using mass flow controllers (Coastal Instruments Inc., Burgaw, North Carolina). The target ozone concentrations were recorded by photometric analyzers (API Model 400, Teledyne, San Diego, California). Mean chamber temperature, relative humidity, and air flow were recorded hourly.

### Glucose tolerance tests (GTT) for all rat experiments

For glucose telemetry, gain of function adrenalectomy experiment, and pharmacological inhibitor studies, GTT was performed in rats after air or ozone exposure as previously described.^18,21,23^ Because rats underwent air or ozone for 4 hours exposure prior to GTT when no food was provided, this served as fasting for GTT. Immediately after air or ozone exposure, baseline blood glucose concentrations (0 hr) were determined by tail prick using a sterile 23-gauge needle with a Bayer Contour Glucometer (Leverkusen, Germany). For glucose telemetry animals, a different glucometer was used (Nova Biomedical StatStrip® Xpress, DSI). Rats were then injected with 20% D-glucose (20% pharmaceutical grade D-glucose; Covetrus, Dublin, OH; diluted to 2 g/kg/10mL, i.p.) as described for telemetry experiment. Glucose levels were measured by tail prick at 30, 60, 90, and 120 min.

### Time course assessment of hormonal response to ozone in rats

A separate cohort of healthy male 12-13-week-old WKY rats was exposed to clean air or ozone at two concentrations (0.4 and 0.8 ppm) for 30 min, 1 hr, 2 hr or 4 hr (n=6-8 per group) and necropsied immediately after each time point (Figure 3A) to collect blood samples for assessment of various pituitary-derived and other hormones. Ozone exposures were performed as described above.

**Figure 1.**
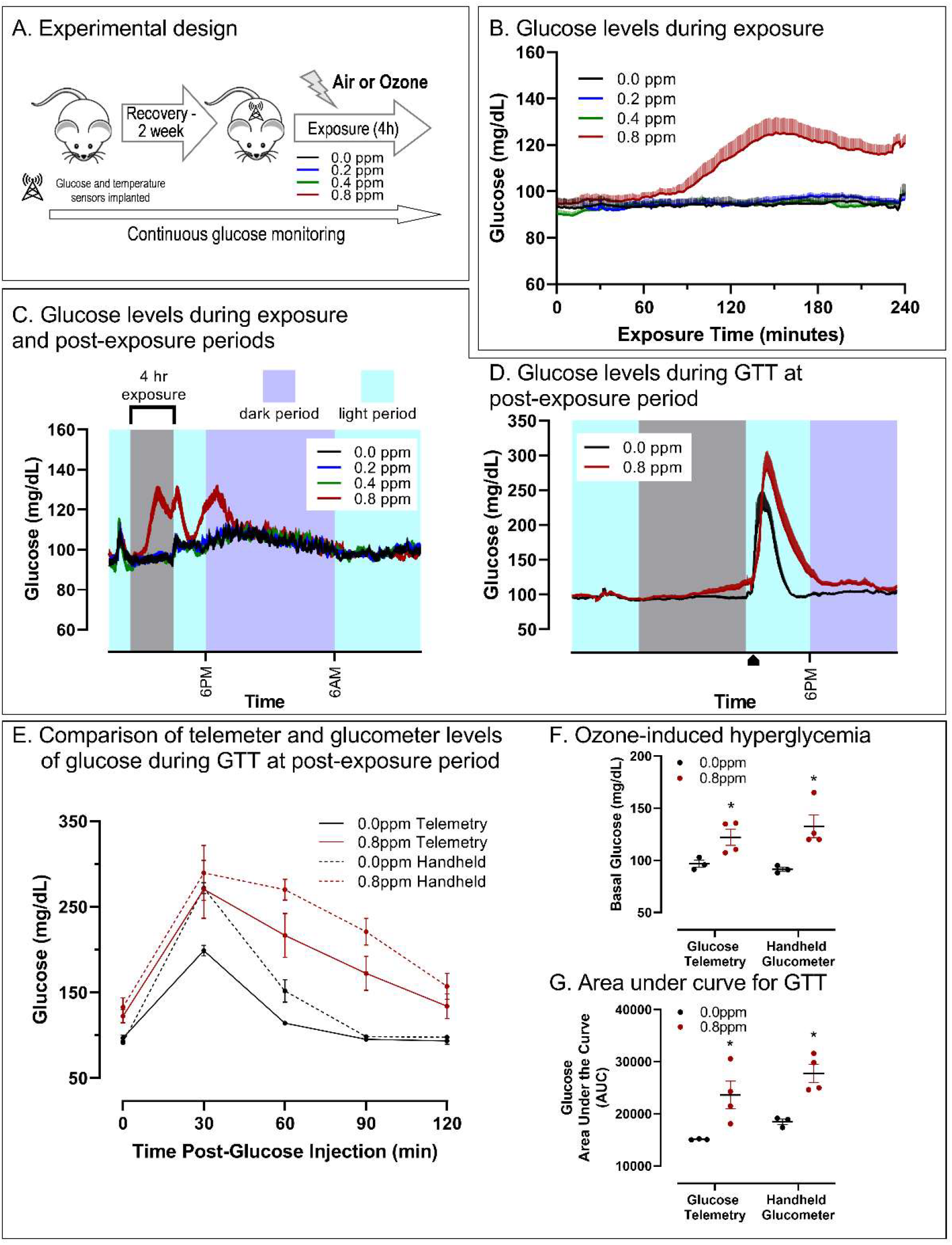
Real-time glucose monitoring in freely moving animals reveal dynamicity of ozone-induced changes and response to glucose injection. A) A schema showing surgical implantation of glucose radiotelemetry devices followed by recovery and then ozone exposure at different concentrations. Glucose levels were measured real-time in a weekly cross over design over 4 weeks and averages of every minute are plotted as mean ± SEM of n=7/concentration. B) Real-time changes in blood glucose levels during 4 hours of air (0.0 ppm) or ozone exposure (0.2, 0.4, and 0.8 ppm). C) Glucose levels during first 24 hours after 4-hour air or ozone exposure (n=7/concentration). D) Glucose levels during 5^th^ and 6^th^ week of cross-over exposure to air or 0.8 ppm ozone and during glucose tolerance test performed immediately following exposure (n=7/concentration). E) Comparison of glucose levels measured through tail prick every 30 min and during continuous monitoring through telemetry (n=7/concentration). F) Comparison of baseline glucose levels assessed using tail prick and telemetry following 4-hour air or ozone exposure. G) Comparison of area under the curve assessment of glucose tolerance test showing similarity in the glucose levels assessed using tail prick and those assessed through telemetry.

**Figure 2.**
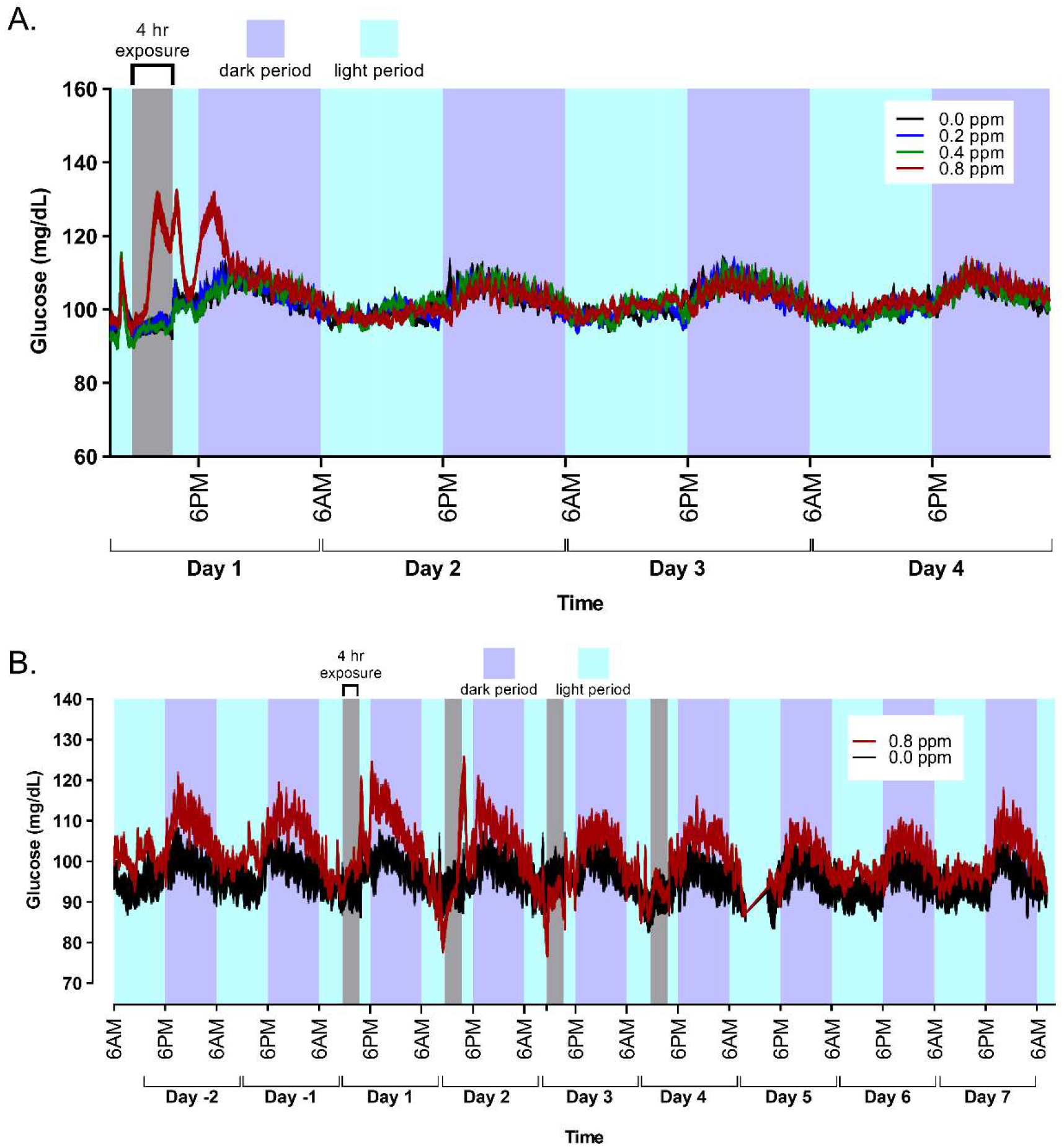
Recovery from a single 4-hour ozone exposure induced hyperglycemia, diurnal variation, and adaptation during repeated ozone exposure. A) Glucose levels were measured real-time in a weekly cross over design over 4 weeks and averaged for every minute are plotted as mean of n=7/concentration. The data show ozone-induced hyperglycemia during and following exposure and reversal of this effect in subsequent non-exposure days. Note the higher levels of circulating glucose at nighttime when animals are active and feeding. B) Ozone-induced hyperglycemia during repeated daily 4-hour exposure for 4 consecutive days followed by 3-day non-exposure period showing nearly complete adaptation on day 3 and day 4 where no ozone effect is evident (n=3-4/group). Data used during telemetry calibration were not used in analysis or graphing.

**Figure 3.**
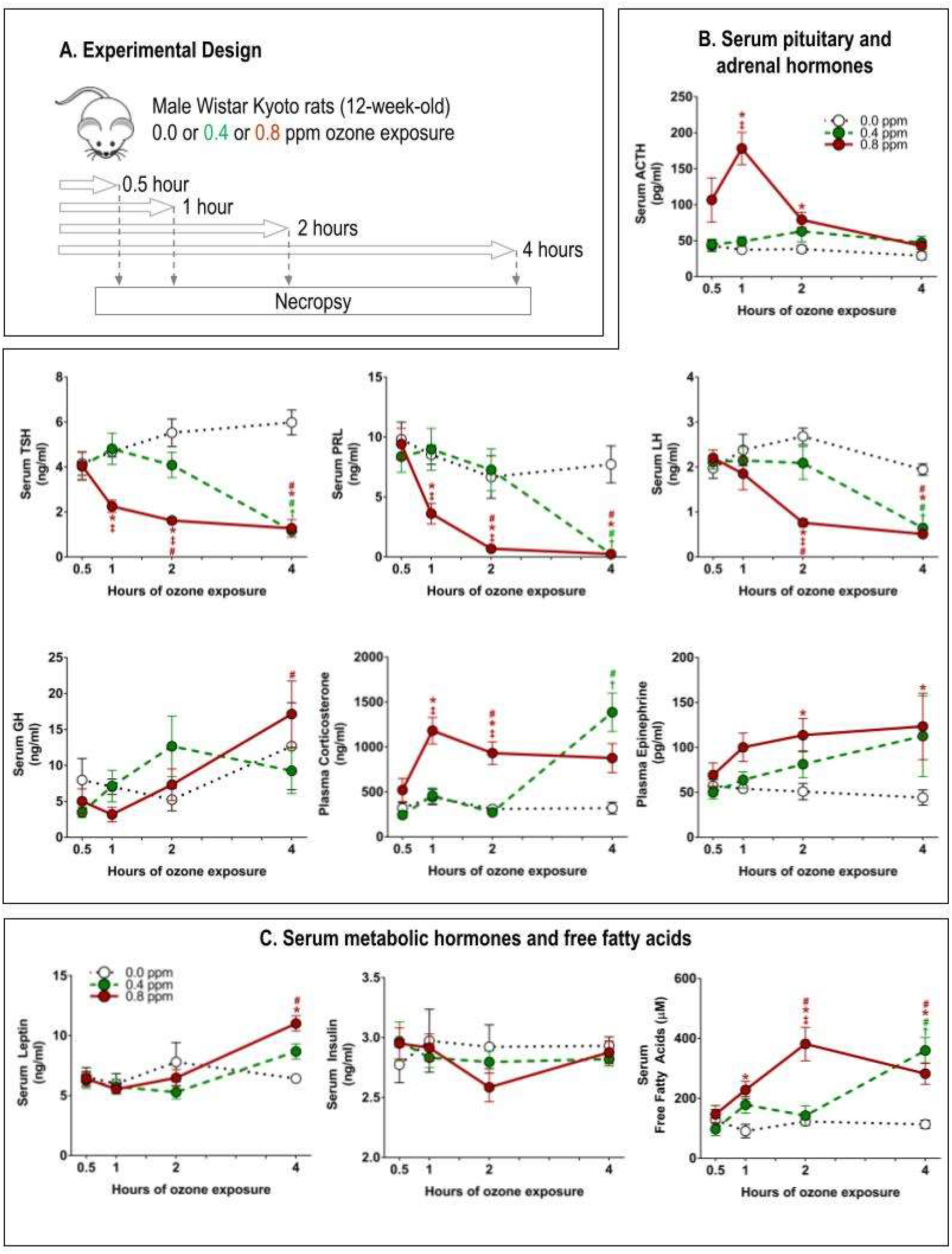
Temporal changes in anterior pituitary, adrenal-derived, and metabolic hormones, as well as circulating metabolites during 4-hour exposure to air or ozone. A separate cohort of healthy rats were exposed to air or ozone at 0.4 or 0.8 ppm for 30 min, 1 hour, 2 hour or 4 hour and necropsies were performed immediately following each exposure (within 15-20 min to collect blood samples). A) experimental design; B) serum pituitary and adrenal-derived stress hormones; and C) metabolic hormones and levels of free fatty acids. Data show mean ± SEM (n=6-8 animals/group). Significant differences (p<0.05) are denoted by “*” for 0.8 ppm versus 0.0 ppm, “†” for 0.4 ppm versus 0.0 ppm, and “‡” for 0.8 ppm versus 0.4 ppm for time matched groups. ACTH, adrenocorticotropic hormone; TSH, thyroid stimulating hormone, PRL, prolactin; LH, luteinizing hormone; GH, growth hormone. Note that the data for corticosterone and epinephrine are recently published in a table form (Henriquez et al., 2021).

### Necropsy and blood samples collection for hormones in time course study

For hormonal time course study, as reported in previous study,^24^ rats were necropsied within 15 minutes after each exposure. Necropsies were performed in a staggered manner. Rats were euthanized with Fatal Plus (sodium pentobarbital, Virbac AH, Inc., Fort Worth, TX; >200 mg/kg, i.p.). Blood samples were collected from the abdominal aorta directly in vacutainer serum separator tubes and EDTA tubes for serum and plasma separation, respectively. Blood samples were spun at 3500 x g for 15 min at 4° C; serum and plasma samples were aliquoted and stored at -80 °C until analysis.

### Plasma and serum analysis of hormones in time course study

Plasma levels of epinephrine (adrenaline) and corticosterone were quantified using kits from Rocky Mountain Diagnostics (Colorado Springs, CO) and Arbor Assays (Ann Arbor, MI), respectively. These two hormones’ data are recently published in a table form along with immune effects.^24^ Serum pituitary hormone levels for adrenocorticotropic hormone (ACTH), thyroid stimulating hormone (TSH), prolactin (PRL), luteinizing hormone (LH), growth hormone (GH), brain-derived neurotrophic factor (BDNF), and follicle stimulating hormone (FSH) were determined using MILLIPLEX MAP Rat Pituitary Magnetic Bead Panel following manufacturer’s protocol (Merck-Millipore, Burlington, MA). Serum free fatty acids were measured using kits from Cell Biolabs, Inc (San Diego, CA) adapted for use on a Konelab Arena 30 clinical analyzer (Thermo Chemical Lab Systems, Espoo, Finland). Serum insulin and leptin were quantified using Meso Scale (Meso Scale Discovery, Gaithersburg, MD).

### Adrenalectomy (AD) and sham (SH) surgeries, drug treatments, and ozone exposure

GTT was performed in our previous study for which pulmonary data are recently published.^25^ In that study, to assess the role of adrenal-derived stress hormones, male WKY rats (12–13 weeks old) that underwent total bilateral adrenalectomy (AD) or control sham (SH) as previously described.^25^Briefly, rats were anesthetized with ketamine (25–50 mg/kg in saline, i.p.), and once anesthetized, injected with the analgesic buprenorphine (0.02 mg/kg/ml in saline, s.c.). During the surgery, anesthesia was maintained by nose-inhalation of vaporized isoflurane (∼3%, 1-2 LPM of O_2_). Animal surgeons from Charles River Laboratories Inc performed the surgeries. Animals were placed in sternal recumbency, and dorsal incision was made. Adrenals from both sides were removed and the muscle layer was sutured to close the abdominal cavity. The surgical wound clips were used to clip the skin and close the wound. SH surgeries were performed using the same anesthesia and surgical approaches as AD except for the removal of adrenal glands. Rats were recovered on heating pads and assessed for signs of distress and pain. Once awake, meloxicam (0.2 mg/kg in saline; s.c.) and buprenorphine (0.02 mg/ml/kg in saline, s.c. every 8–12h for 2 times) was administered for analgesia. After the surgery, AD rats received water with 0.9% NaCl to maintain adequate salt-water balance in the absence of mineralocorticoids eliminated due to AD along with other adrenal-derived stress hormones. All animals were provided with powdered as well as pelleted food *ad libitum*. The rats were pair housed with EnviroDry enrichment/nesting material and allowed to recover for 4–6 days prior to drug treatment as reported in our previous companion paper^25^ and described below. These SH and AD rats, as detailed in our recent publication^25^, were randomized by body weight into four groups (vehicle: air, vehicle: ozone, clenbuterol (CLEN; β2AR agonist) plus dexamethasone (DEX; GR agonist):air, and CLEN+DEX: ozone) resulting in 8 total groups (n=8/group). In brief, after 4-6 days of recovery from SH and AD, rats were treated with vehicles (saline 1 mL/kg as control for CLEN, i.p.) and corn oil (1 mL/kg as control for DEX, s.c.) or clenbuterol hydrochloride, a long acting β2AR agonist (CLEN; 0.2 mg/kg in saline, i.p.) and GR agonist, dexamethasone (DEX; 2 mg/ml corn oil/kg, s.c.). Generally, CLEN injections were followed by DEX. The drug treatment began 1 day prior to start of ozone exposure and continued the day of air or ozone exposure in the morning at ∼6am. These high doses were selected to restore the depleted activities of epinephrine and corticosterone in AD groups. CLEN and DEX doses are comparable to those used in other controlled experiments using rodents and are sufficient to induce bronchodilation and immunosuppression, respectively.^26,27^ These rats were exposed to air or 0.8 ppm ozone for 4 hours and GTT were performed immediately after exposure as described above.

### Treatment of rats with βAR and GR antagonists and ozone exposure

GTT assessment was also performed in three additional independent experiments, which were carried out to evaluate the role of 1) β adrenergic receptor antagonist propranolol (PROP), 2) glucocorticoid receptor antagonist mifepristone (MIFE) individually, or 3) both in combination to determine their influence on ozone pulmonary effects as published previously.^21^ For each study, 12-13-week-old male WKY rats were randomized by body weight into four groups (vehicle/air, drug/air, vehicle/ozone, drug/ozone, n = 8/group). In the first experiment, rats were injected with either sterile saline (vehicle; 1 mL/kg, i.p.) or propranolol hydrochloride (PROP, Sigma-Aldrich, St Louis, MO; 10 mg/kg in saline, i.p.). In the second experiment, rats were injected with pharmaceutical grade corn oil (vehicle; 1 mL/kg, s.c.) or mifepristone (MIFE, Cayman Chemical Co., Ann Arbor, MI; 30 mg/kg in corn oil, s.c.). In the third experiment, rats were injected with vehicles, saline (1 ml/kg, i.p.) followed by corn oil (1 ml/kg, s.c.) or drugs, PROP (10 mg/kg, i.p.) followed by MIFE (30 mg/kg, s.c.) (PROP+MIFE). The rationale for drug selection and treatment protocol are explained in our previous companion paper describing pulmonary and immune responses.^21^ To assure complete inhibition of βAR and GR receptors, the daily morning treatment began 7 days prior to the air or ozone exposure and was continued the day of exposure. In each study, rats were exposed to filtered air or 0.8 ppm ozone for 4 hours as indicated above, and GTT was performed immediately after exposure.

### Assessment of glucose and cortisol in human clinical study samples

Human plasma samples were obtained from a clinical study conducted through the University of North Carolina (Chapel Hill, NC) under IRB# #13-1644. The study involved exposure of young healthy human volunteers to filtered air or 0.3 ppm ozone exposure in a cross-over design where the same subjects were randomly exposed to air or ozone during two distinct visits that were separated by two weeks or longer. Prior consents were obtained from all individuals participating in the study. Blood samples were collected prior to and immediately following 2-hour exposure to air or ozone for each of 34 subjects. Demographic information is provided in Table S2. For human plasma samples, glucose levels were analyzed using Bayer Contour Glucometer and test strips (Leverkusen, Germany). Plasma cortisol levels were analyzed using human cortisol kit from Arbor Assays (Ann Arbor, MI).

### Statistics

For all analyzed endpoints, a threshold of p < .05 was used to determine significant effects. Outliers were identified using the boxplot method, defined as above Q3 + 1.5 IQR or below Q1 – 1.5 IQR and discarded. Analysis of glucose telemetry data was done by assessing peak glucose levels and area under the curve (AUC) for set timepoints (i.e. during exposure, during GTT, etc.). AUC was calculated using the trapezoidal method as previously described.^22^ Significant effects of exposure were determined using one-way analysis of variance (ANOVA) with a Tukey’s post hoc comparison to assess differences between ozone concentrations or a student’s t-test when only one ozone concentration was used. To determine the relationship between glucose and animal body temperature, a linear regression was performed. To compare the similarities of glucose telemetry and handheld glucometer, significant effects were examined using a two-way ANOVA (measurement technique, exposure) with a Tukey’s post hoc comparison used for pair-wise comparisons. To determine adaptation over time, a two-way repeated measures ANOVA was performed (exposure, day) with Tukey’s post-hoc comparison used for pair-wise comparisons. For analysis of the effects of ozone on serum pituitary and adrenal hormones and serum metabolic hormones and free fatty acids, a two-way repeated measures ANOVA (exposure, time) was performed and followed by Tukey’s post-hoc comparisons for each ozone concentration. To explore the effects of adrenalectomy and/or pharmaceutical interventions on ozone’s glucose-inducing effects, a three-way (exposure, drug, surgery) or two-way (exposure, drug) ANOVA was performed with Tukey’s post hoc used for pairwise comparisons. For human data, the percent change in plasma levels of glucose and cortisol pre- and post-exposure were calculated and used for paired-samples t-test. GraphPad Prism 9 (version 9.1.2) was used for statistical analysis and graph generation. Statistical summaries pertaining to all data presented in figures is provided the Supplemental Excel File (Excel Table S1) with a detailed breakdown of all statistical analyses, effect size calculations, and notes on any outliers removed.

## RESULTS

### Real-time *in vivo* glucose monitoring during and after ozone exposure

We used a novel real-time blood glucose telemetry system (Figure 1A and Figure S1)^20^ that has not been employed in previous air pollution studies. Using a cross-over design with the 7 telemetered rats, we obtained independent readings at each concentration for a weekly 4-hour exposure to air or ozone with 1-week washout (Table S1). We have shown that one week wash-out period after a single 4-hour ozone exposure is sufficient to clear effects from a previous exposure in rats.^28^ Continuous monitoring of glucose during 4 hours of ozone exposure at various concentrations and post exposure periods in rats allowed insights in precise timing for a stressor to impact changes in circulating glucose and the longevity of a stress response. It also allowed monitoring of glucose intake-related and diurnal changes. Because glucose telemetry also included the assessment of core body temperature, we were able to show that a drop in core body temperature correlated with glucose changes.

Hyperglycemia began to occur at ∼90 min into a 4-hour ozone exposure but only in the 0.8 ppm group (Figure 1B). In the 0.8 ppm group, both peak glucose and AUC are significantly increased, and glucose plateaued between 2.5 and 3 hours, followed by a second peak between 4-5 hours during the first day of exposure. These changes may reflect fine oscillatory adjustment in homeostatic processes, which might be important for centrally balanced and precisely controlled responses to ozone stress. A third peak of hyperglycemia in the 0.8 ppm ozone group was noted roughly 1 hour after the beginning of dark cycle (and 4 hours post cessation of ozone exposure) when rodents were active and feeding (Figure 1C). The increase in glucose at 90-min was strongly associated with hypothermia in the 0.8 ppm ozone group, but without the fluctuations seen in glucose levels (Figure S2).

After completing 4-week exposure using cross over design, animals were assigned air or 0.8 ppm ozone group for subsequent weeks. On the 5^th^ and 6^th^ week, rats were exposed to air or 0.8 ppm ozone for 4 hours using crossover design (alternating exposure assignment to air or ozone each week), and GTT was performed immediately following exposure in telemetered rats. These rats exhibited ozone-induced glucose intolerance in addition to hyperglycemia, as evidenced by both increased peak glucose levels and increased AUC (Figure 1D) consistent with our previous studies involving post exposure assessment.^18^ The telemetry data for blood glucose after bolus glucose injection matched the data obtained through a handheld glucometer (Figure 1E), with both telemetry and glucometer finding a significant increase in basal glucose (Figure 1F) and AUC (Figure 1G) during the 0.8 ppm exposure and no difference between measurement techniques (glucometer vs telemetry). Combined, these data suggest that glucose monitoring in real-time offers opportunities to concurrently assess effects of dietary glucose and acute environmental stressor exposure. The temporal co-occurrence of ozone-induced hypothermia and hyperglycemia in the 0.8 ppm group suggests that these processes are linked or induced through common upstream events (Figure S3A). This hypothermia did not occur after glucose injection. In general, when glucose levels are high, core body temperature increases along with activity as noted during the dark cycle for rodents (Figure S3B). However, ozone-induced changes in blood glucose and body temperature were in the opposite direction, suggesting stress-induced disturbance in homeostasis to conserve metabolic energy and direct it where needed. Hypothermia, which has been linked to stress-induced glucocorticoid increases in humans^29^ was also evident after ozone exposure in our previous study that employed real-time ECG monitoring in rats.^30^

### Real-time glucose monitoring during ozone adaptation and diurnal variation

After a single 4-hour 0.8 ppm ozone exposure, hyperglycemia was not noted during subsequent days of no exposure (Figure 2A). The diurnal changes were apparent in all animals showing higher levels of glucose at nighttime when compared to daytime. Repeated daily exposure to ozone has been associated with adaptation/tolerance in mice.^31^ However, the mechanism of adaptation remains elusive. With the use of glucose telemetry, we linked ozone adaptation to glucose changes and obtained precise timing of adaptation. On week 7, animals were exposed to air or 0.8 ppm ozone (4 hours each day) for 4 consecutive days to determine if adaptation occurs with regards to glucose and body temperature changes during continued daily exposure. Because of the repeated exposures during week 7 (daily for 4 days), crossover design was not possible and therefore sample sizes are smaller than for all other experiments (air, n=3; ozone, n=4). Ozone at 0.8 ppm 4 hours/day for 4 consecutive days led to significant increases in peak glucose and glucose AUC only on days 1 and 2, with a notable lack of increase in blood glucose during and right after the exposure on the third and fourth days despite continued exposure(Figure 2B). Near complete adaptation to ozone exposure was evident by the 3^rd^ day despite continued exposure on 3^rd^ and 4^th^ day. Thus, continuous glucose monitoring indicated that this adaptation occurs on the 3^rd^ day but not on the 2^nd^ day of ozone exposure, and is associated with attenuation of ozone-induced hyperglycemia (Figure 2B). The adaptation was also noted in hypothermia on the 3^rd^ day (Figure S3C). These ozone-induced changes in circulating glucose reflect the status of glucose metabolic processes in tissues.^28^ Thus, real-time glucose monitoring allows one to assess the timings of metabolic alterations and adaptation during ozone exposure.

### Temporality of ozone-induced HPA and SAM activation

Because the method for continuous monitoring of corticosterone in animals is still not available, we exposed a separate cohort of naïve rats to air or ozone for variable durations spanning a 4-hour time frame to assess temporal changes in adrenal-derived and other neuroendocrine hormones (Figure 3A). This study followed a similar paradigm to real-time glucose monitoring but used a distinct cohort of rats at each timepoint during 4-hour ozone exposure to assess serum samples for key pituitary, adrenal-derived, and metabolic hormones (Figure 3). A sharp rise in adrenocorticotropic hormone (ACTH) occurred at 30 min into the 0.8 ppm ozone exposure and peaked at 1 hour, reflecting the activation of the HPA axis and concomitant ACTH release from the anterior pituitary as early as 30 min (Figure 3B). Upon receiving stress signals in the paraventricular nucleus of the hypothalamus, the secreted corticotrophin releasing hormone traverses to the pituitary through the hypothalamic-pituitary portal system and activates ACTH secretion from the anterior pituitary. ACTH released into the systemic circulation reaches the adrenal cortex to stimulate corticosterone/cortisol synthesis and release involving hypothalamus-pituitary-adrenal (HPA) axis.^10^ Consistent with this, we noted that the levels of circulating corticosterone in rats increased starting at 1 hour into ozone exposure (0.8 ppm) prior to the increase in glucose and remained significantly elevated until 4 hours of exposure (Figure 3C)^24^ despite the restoration of ACTH to baseline levels at this time point. It is important to note that although no significant ACTH increase occurred during the 0.4 ppm ozone exposure, the increase in corticosterone was significant at 4 hours, suggesting that the stress response was concentration dependent. The data for corticosterone changes were recently published in a table form.^24^

The stress response induced by acute physical and emotional stress also involves the activation of the splanchnic sympathetic nerves via the hypothalamus leading to a release of epinephrine from chromaffin cells of the adrenal medulla.^32^ The sympathetically-mediated (SAM axis) epinephrine increase generally precedes the increases in corticosterone during a fight-or-flight response. The data show that after 30 min of 0.8 ppm ozone exposure, the levels of epinephrine were increased as recently reported.^32^ Exposure to the lower concentration of ozone (0.4 ppm) also caused an increase in epinephrine, although the time required for this increase was longer, as it was evident only after 4 hours of exposure (Figure 3B).^32^ The sustained increases in circulating epinephrine during the 4 hour of ozone exposure (Figure 2B) also corroborated our prior findings involving a single end of exposure measurement.^18,23^

### Temporal changes in HPT and HPG hormones during ozone exposure

Because ozone-mediated stimulation of the neuroendocrine system may also influence other hypothalamic stress pathways such as hypothalamic-pituitary-thyroid (HPT) and hypothalamic-pituitary-gonadal (HPG) axes^33^, we next assessed temporal effects of ozone exposure on relevant hormones. We demonstrate here that ozone exposure results in time- and concentration-dependent decline in TSH levels, which occurs sooner (1 hour) at 0.8 ppm and is temporally linked to increases in corticosterone and precedes the increased glucose response with 0.8 ppm ozone (Figure 3B).We also assessed pituitary hormones involved in gonadal axis includin follicle stimulating hormone (FSH), prolactin (PRL), and luteinizing hormone (LH) (Figure 3B). We report that ozone exposure depleted not only LH, but also PRL (>95%), with the 0.8 ppm concentration causing a more rapid depletion. This corroborates our recent study where we reported a decrease in circulating PRL assessed once after 4 hour of ozone exposure.^33^ Here, the temporal assessment shows depletion of circulating PRL as early as 1 hour into exposure concomitant with peak ACTH levels. However, LH levels did not decrease until 2-hour of exposure (Figure 3B) and the levels of FSH were not changed after ozone exposure in male WKY rats (*data not shown*). These data provide insights on how acute ozone inhalation can dynamically and differentially impact various neuroendocrine axes that have major impact on homeostatic physiological processes.

Because pituitary-derived growth hormone (GH) is involved in cell growth and regeneration processes,^34^ and tied to metabolic changes,^35^ we assessed the kinetics of growth hormone changes and noted that a delayed but concentration-dependent increase in GH occurred at 4-hour after ozone exposure suggesting that the anabolic processes are being activated (Figure 3B). A similar temporal pattern was noted for the increase in leptin, which regulates satiety at the level of hypothalamus (Figure 3C). On the other hand, the increase in circulating free fatty acids was noted as early as 1 hour after ozone exposure, suggesting the early activation of lipolytic activity in adipose tissue coinciding with changes in circulating adrenal-derived hormones, but a delayed increase in GH coincides with leptin release from adipose tissue (Figure 3C). Overall, ozone-induced hyperglycemia is reflective of neuroendocrine responses that might involve a wide array of changes in metabolic and cell growth processes.

### The role of epinephrine and corticosterone in mediating hyperglycemia

Adrenal-derived epinephrine and glucocorticoids are the major regulators of liver metabolic processes during stress,^36^ and adrenalectomy diminishes ozone-induced hyperglycemia and glucose intolerance.^22^ Therefore, we further assessed the roles of adrenal-derived stress hormones in mediating changes in circulating glucose after ozone exposure. Ozone-induced hyperglycemia and glucose intolerance were nearly eliminated in vehicle-treated AD rats, confirming our earlier findings.^22^ Moreover, all animals treated with CLEN+DEX developed marked hyperglycemia and glucose intolerance, in both SH and AD rats exposed to air. Further, this response was exacerbated in rats exposed to ozone demonstrating βAR+GR activation through increased epinephrine and corticosterone mediating hyperglycemia (Figure 4B). PROP or MIFE given individually did not reduce ozone-induced hyperglycemia, but PROP+MIFE in combination significantly decreased hyperglycemia severity. Similarly, the blockade of PROP and MIFE individually only partially reversed ozone-induced glucose intolerance, but together, they markedly diminished ozone-induced glucose intolerance, suggesting the involvement of both epinephrine and glucocorticoids in mediating glucose increases during ozone exposure (Figure 4C). These results indicate a causal link between ozone-induced increases in the release of adrenal-derived stress hormones through SAM and HPA activation and resultant glucose metabolic alterations. Together, these data suggest that there are dynamic relationships between ozone-induced changes in neuroendocrine stress pathways, HPA activation and glucose homeostatic changes.

**Figure 4.**
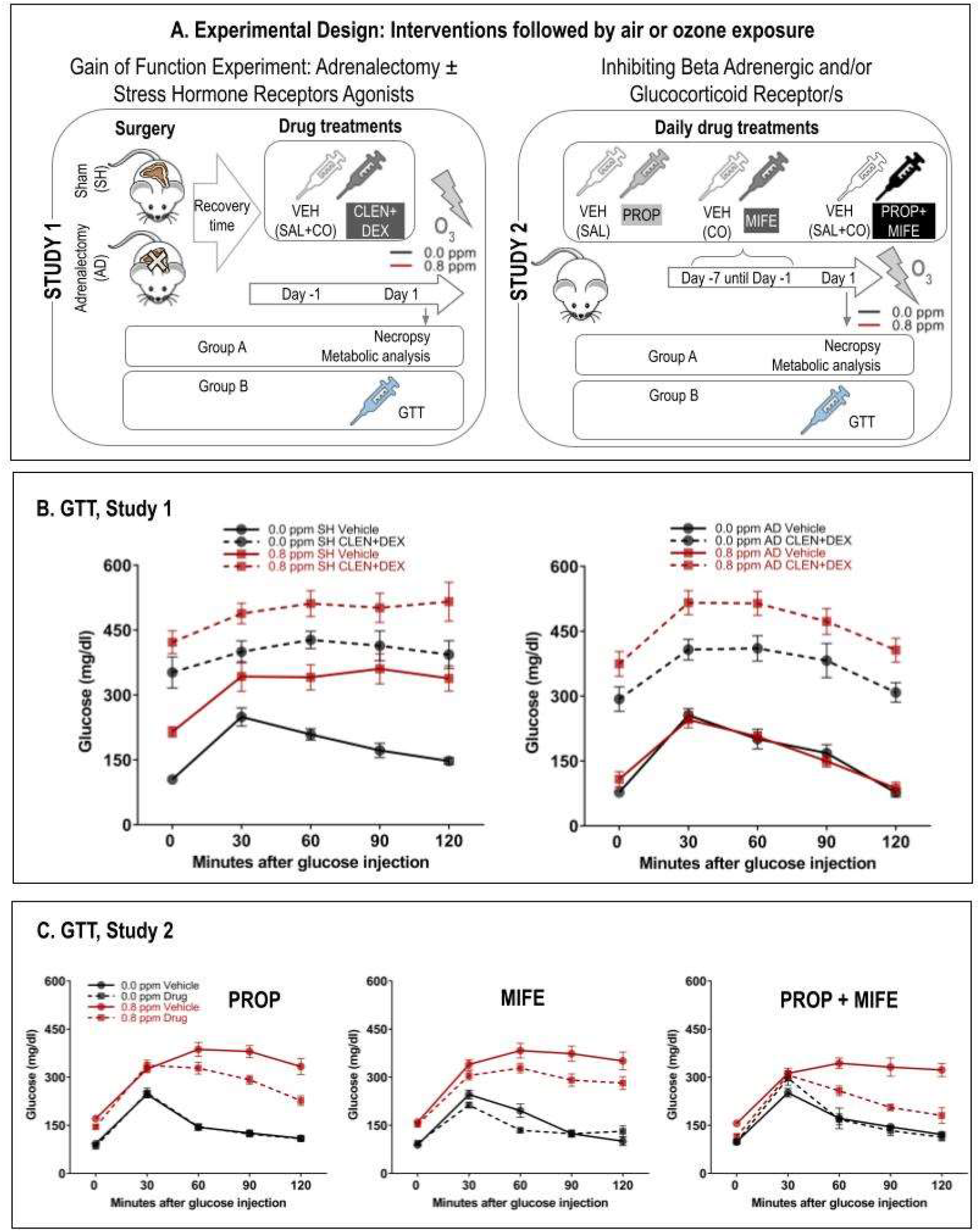
Mechanistic link between blood glucose changes and adrenal-derived stress hormones as determined using adrenalectomy (AD), and pharmacological interventions. A. Left panel shows the experimental design involving AD and treatment of rats with vehicle (VEH) or β2AR plus GR agonists (clenbuterol [CLEN] + dexamethasone [DEX]) 1-day prior to and the day of air or ozone exposure (Henriquez et al., 2018a). Right panel shows the treatment of healthy rats with vehicle (VEH) or βAR and/or GR blockers (propranolol [PROP] and mifepristone [MIFE], respectively, followed by air or ozone exposure (Henriquez et al., 2018b). These published papers evaluated pulmonary effects of interventions. To assure effective receptor blockade the treatment began 7 days prior to air or ozone exposure. B and C. Glucose tolerance test (GTT) and the glucose data at baseline and following glucose injection in each study (n=6-8). SH, sham surgery; AD, adrenalectomy surgery; VEH, vehicle; CLEN, clenbuterol; DEX, dexamethasone; PROP, propranolol; MIFE, mifepristone; SAL, saline; CO, corn oil.

### Ozone-induced stress response in humans

Hormonal response to stress is conserved across species; in this study, we wanted to determine if the results seen in rats would extend to humans. We assessed glucose and cortisol levels in plasma samples of human volunteers exposed to air or 0.3 ppm ozone during 2 hours of intermittent exercise in a cross-over design where each volunteer was exposed to air and ozone but separated by at least two weeks interval (IRB# #13-1644; Figure 5A). There were no differences in blood glucose levels between volunteers exposed to air or ozone (Figure 5B), likely because all subjects were exposed in a protocol involving intermittent exercise. In contrast, although exercise resulted in decreased cortisol levels in air exposed individuals, this decrease was significantly attenuated in volunteers exposed to ozone as determined by % change between pre- and postexposure (Figure 5B).

**Figure 5.**
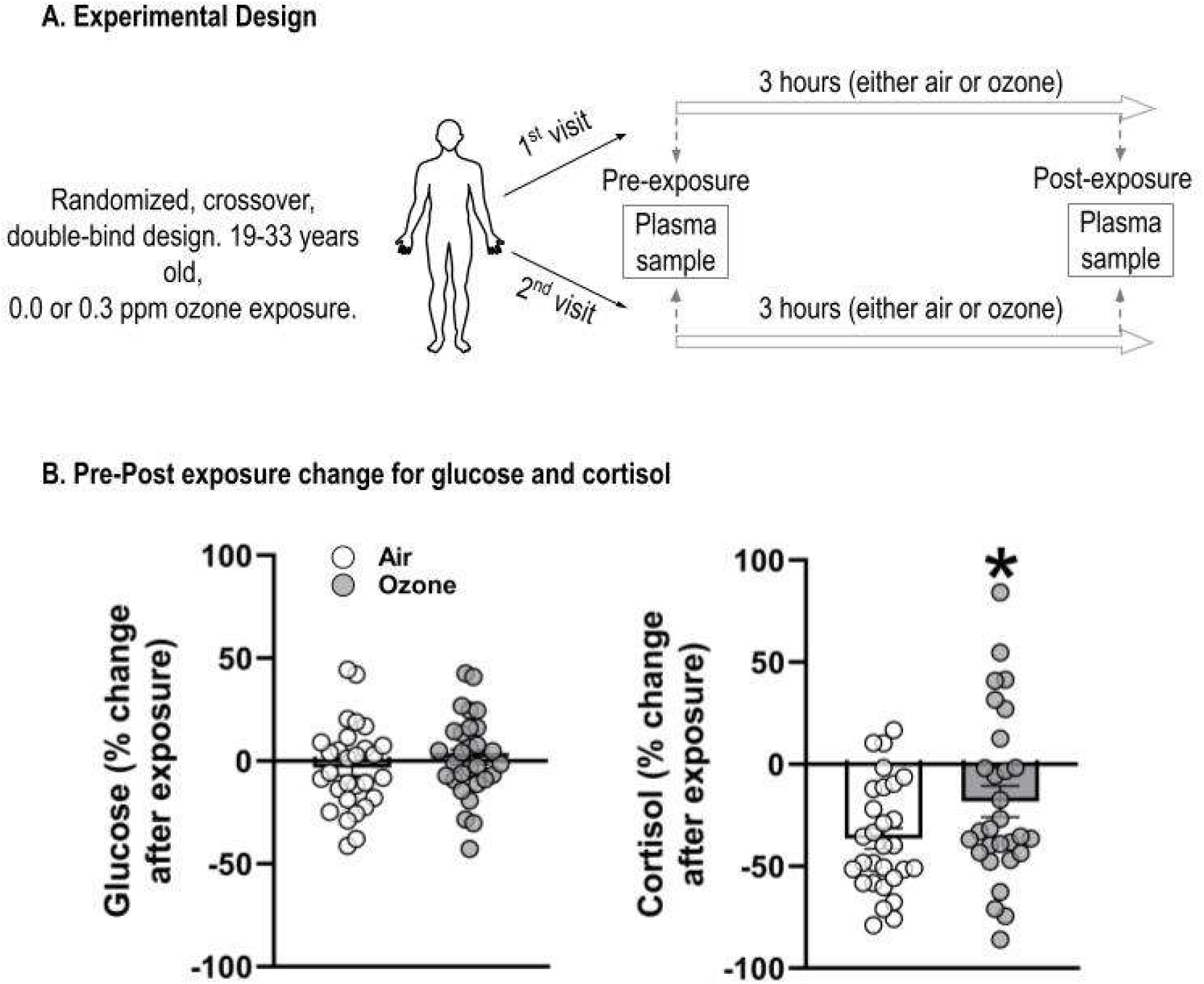
Changes in plasma glucose and cortisol in humans within 1-2 hour following exposure to air or 0.3 ppm ozone. (A) Archived serum samples from a clinical study were obtained where blood samples were collected pre and post exposure during two clinical visits from healthy young volunteers who are exposed to air or 0.3 ppm ozone for 2 hours in a cross-over design separated for at least 2 weeks. During air or ozone exposure volunteers were intermittently exercising. B. There was no significant increase in blood glucose likely due to exercise, however, normal cortisol decreases from exercise were significantly attenuated (paired-samples t-test; n=29; p = 0.02). Samples > 1.5 interquartile range were removed as outliers.

## DISCUSSION

Prior research on air pollution health effects focused primarily on cardiorespiratory outcomes elicited by local effects in the lung. Recently it has become apparent that inhaled irritant air pollutants might be perceived as stressors by the autonomic nervous system, resulting in stimulation of neuroendocrine axes which mediate acute effects in the lung and periphery.^37^ The impairment and/or persistent hyperactivity of stress responses has been linked to chronic psychiatric, neurobehavioral, cardiometabolic, and reproductive health abnormalities. Because stress responses are dynamic and CNS-regulated to induce reversible peripheral changes, its real-time assessment is necessary for in-depth understanding of its impact on health and resiliency. We used real-time glucose telemetry in rats, combined with temporal assessment of neuroendocrine hormones to demonstrate in rodents a contextual relationship between air pollution induced stress response and hyperglycemia. We demonstrated a mechanistic link between hyperglycemia and adrenal-derived stress hormones using adrenalectomy and pharmacological interventions of adrenergic and glucocorticoid receptors (βAR and GR). Further, we showed that exercise-induced depletion of cortisol in humans was significantly attenuated by ozone, supporting the role of HPA axis consistent with our prior human study.^19^ Thus, real-time monitoring of glucose and stress hormones may serve as immediate biomarkers of air-pollutant-induced pulmonary stress and the interactive impacts of other environmental contaminants and non-chemical stressors.

Hyperglycemia is one of the earliest markers of stress-induced homeostatic changes that is consistently noted after a single ozone exposure^18,22,23^ and other irritant exposures.^38^ Here, we show that the temporality of this neuroendocrine response induced by ozone can be assessed using real-time glucose telemetry in rats. Upon daily ozone exposure for 3 or more consecutive days, the ozone-induced hyperglycemic and hypothermic response is no longer evident, reflective of stress adaptation. We show that ozone-induced hyperglycemia, which coincides with hypothermia during exposure, is preceded by changes in circulating anterior pituitary-derived hormones and the release of adrenal-derived epinephrine and corticosterone, linking the activation of SAM and HPA axes and the inhibition of HPT and HPG axes to hyperglycemia. Further, eliminating adrenal-derived stress hormones from circulation through adrenalectomy or pharmacologically inhibition of stress hormone receptors diminishes ozone-induced hyperglycemia, and treatment with stress hormone receptors agonists restores ozone-induced hyperglycemia in adrenalectomized rats. Combined, these results highlight the utility of real-time glucose measurement as a sensitive marker that reflects the impacts of air-pollutant exposure, and may be useful in linking source to health outcomes.

Acute ozone exposure has been shown to activate stress-responsive regions in the rat brain.^39^ The release of stress hormones by activation of SAM and HPA axes is integral in mediating key metabolic processes that channel energy resources where needed by acting on AR and GR.^40^ Glucocorticoid feedback regulation on HPA activity, and the roles of mineralocorticoids and catecholamines, have been implicated in stress adaptation and the plasticity of organismal neural responses.^10^ However, the full understanding of molecular mechanisms linked to impaired stress adaptation and neuropsychiatric disorders is still lacking. The adaptation from ozone-induced hyperglycemia on the third consecutive day of exposure suggests a neuroendocrine contribution to ozone adaptation. This adaptation response was not evident on the second day of ozone exposure. Moreover, one-week of no exposure washout-period was associated with the loss of adaptation, indicating remarkable plasticity that may be influenced by stressor type, potency, and longevity of exposure. Given the contribution of glucocorticoids in stress adaptation,^11^ and the link between changes in circulating cortisol and chronic neurobehavioral and metabolic diseases,^41,42^ the temporal assessment of changes in glucose and cortisol is critical for evaluation of health status, resiliency, and longevity of stressor effects. Further, the loss of dynamicity or oscillatory rhythms is an important indicator of chronic health issues,^43^ which may provide additional insight from temporal monitoring.

Real-time glucose monitoring is now more frequently employed for diabetic individuals who require repeated assessment of blood glucose. In our study, we were able to assess increases in blood glucose during GTT and from food intake during nighttime in rats. Thus, the real-time glucose assessment offers the opportunity to study interactive effects of diet, metabolic disease, and stress from air pollutant exposures. Stress in diabetics may exacerbate preexistent hyperglycemia as reported in diabetic Goto Kakizaki rats^44^ and in diabetic patients.^45^ Thus, the temporal assessment of glucose could provide insights on influence of stressors on overall physiological status and underlying metabolic health condition.

Combined, evidence of temporal and concentration-dependent changes in hormones associated with not only SAM and HPA, but also HPT and HPG axes during acute ozone exposure, indicate that the stress response involves a complex interplay of multiple neuroendocrine pathways. The selective activation of HPA and SAM axes was associated with concurrent inhibition of HPT and HPG axes as observed by changes in respective hormones. These responses are consistent with acute psychosocial stress-induced increases in ACTH and cortisol, which are associated with depletion of gonadal hormones in men and women.^46^ This is in contrast, however, with concurrent increases in thyroid hormone and corticosterone after exercise in rats.^47^ With regards to HPG hormones, a rapid decline PRL in this study and the reported reversal of its depletion in adrenalectomized rats^33^ suggests the possible involvement of glucocorticoid feedback regulation. While circulating glucocorticoids might also inhibit LH secretion after ozone exposure, a role of gonadotropin releasing hormone and gonadotropin inhibitory hormone is likely in ozone-induced inhibition of LH.^48^ These findings indicate that stress responses may not be uniform between stressor types and that ozone may impact specific neuroendocrine pathways that involve input from multiple interactive signaling processes in the brain to develop a tailored, integrated and temporally regulated host response.

Circulating corticosterone, cytokines, exhausting exercise, caloric deprivation, and even sepsis are linked to depletion of TSH, T4, and T3 in a stress paradigm.^49^ Acute exposure to ozone has been shown to deplete circulating thyroxine and THS in rats.^50^ We have shown that AD reverses ozone-induced inhibition of TSH release, suggesting a role for circulating adrenal-derived stress hormones.^33^ Pituitary TSH release is regulated by hypothalamic thyrotropin releasing hormone (TRH) with feedback controls on HPT at different levels.^51^ Thus, stressor specific differences in activation versus inhibition of given hormonal systems may impact downstream physiological responses. The mechanisms by which anabolic pituitary hormones, such as ACTH and GH, increase whereas the catabolic hormones, such as TSH, FSH, LH and PRL, decrease after ozone exposure may involve a precise and selective activation or inhibition of upstream regulators of hormonal responses after ozone exposure, such as the activation of CRH neurons, FK506 binding protein regulation of glucocorticoid feedback, and other neurotropic factors within the stress-responsive regions impacted by ozone.^10,39^

We have previously noted that ozone-induces pulmonary and liver transcriptional changes reflective of processes that regulate metabolism, cell cycle and regeneration.^52,53^ Because pituitary-derived GH is involved in these processes,^34^ and tied to metabolic changes,^35^ we assessed kinetics of GH changes and noted a delayed but concentration-dependent increase at 4-hour after ozone exposure, suggesting that anabolic processes are being activated. Similar patterns were noted for increases in leptin, which regulates satiety at the level of the hypothalamus. However, the increase in circulating free fatty acids was noted as early as 1 hour, suggesting early activation of lipolytic activity in adipose tissue coinciding with changes in circulating adrenal-derived hormones, but a delayed increase in GH coincides with leptin release from adipose tissue.

Our approach focused on establishing a causal role of SAM and HPA axes activation on hyperglycemia response in a gain of function experiment using AD and stress hormone receptor agonists/antagonists.^25^ Increased circulating epinephrine and glucocorticoids regulate glucose metabolic processes, in addition to regulating other physiological and immunological processes through their action on multiple tissues, including the liver and pancreas.^54^ AD or treatment with pharmacological blockers of βAR plus GR inhibited ozone-induced hyperglycemia and glucose intolerance whereas the combination of βAR and GR agonists amplified ozone-induced hyperglycemia and glucose intolerance. We have previously shown that ozone exposure is associated with increased gluconeogenesis and inhibition of glucose-mediated insulin secretion in rats.^28^ Each AR and GR subtype may be selectively influencing different processes of glucose metabolism, such as gluconeogenesis, and β-cell insulin secretion in the liver and pancreas. These changes following acute ozone exposure suggest that adrenal-derived hormones regulate glucose homeostatic changes induced by ozone.

Stress response is proportional to stressor severity and duration, and is precisely directed to the affected organ system. Furthermore, this response is reversible upon stress discontinuation and in some cases, even after continued stressor application (habituation). However, these reversible physiological stressor effects in healthy individuals can be impaired in susceptible individuals both at the CNS and peripheral organ levels, contributing to health burdens from environmental exposure.^43^ Based on evidence presented in this paper, we surmise that evaluation of the dynamicity of this response through real-time glucose and cortisol monitoring could unravel critical information on the magnitude and persistence of stress from environmental exposures and its impairment in individuals with preexisting diseases including psychosocial and metabolic. Using ozone inhalation as an example, we here demonstrated the utility of such an approach to monitor the dynamics of stressor effects on health that is amenable in humans with currently available technologies.^55^

This study assessed responses after exposure to only ozone. Whereas we have shown similar changes in stress hormones and glucose after exposure to other gaseous irritant, acrolein,^38^ this response may be linked to irritancy and should not be generalized to all pollutants and pollution mixtures. Similarly, the nature and timing of responses to other stressors may vary leading to differences in the organ being affected and the pathology. Moreover, this study assessed only acute health outcomes in healthy animals after inhaled ozone exposure; however, the mechanisms by which repeated exposures and underlying health conditions increase disease susceptibility can be better studied using models of compromised health status. Finally, variations between humans and rodents in respiratory anatomy and physiology as well the uniqueness of the hypothermic response to ozone in rats are important caveats when translating findings to humans. The influence of physiological status at the time of stress encounter is also critical, for example, an observed lack of increase in glucose, but evidence of cortisol response after ozone exposure in humans during intermittent exercise.

In conclusion, we show that health effects of air pollution-induced stress can be monitored in real-time by assessment of blood glucose using telemetry and where possible cortisol/corticosterone in experimental models and in humans. Our data show that exposure to prototypic air pollutant (ozone) rapidly induces anabolic (ACTH, GH) neuroendocrine pathways involving sympathetic activation and adrenal medullary release of epinephrine and HPA-mediated release of glucocorticoids, while inhibiting catabolic (HPT, HPG) pathways. These reversible and dynamic neuroendocrine changes are followed by systemic metabolic alterations and hyperglycemia as well as hypothermia during ozone exposure. The use of real-time glucose telemetry is rats provided insights into the precise temporal relation to changes in stress hormones. We further mechanistically confirm the contribution of adrenal-derived stress hormones based on the evidence that AD and pharmacological inhibitors of stress hormones eliminate, and agonists reverse ozone-induced hyperglycemia and glucose intolerance. Because dynamic changes in circulating stress hormones likely mediate interactive metabolic effects of irritant air pollutants such as ozone, this approach of continuous monitoring of glucose and stress hormones may be useful in assessing health effects and susceptibility variations in epidemiological studies.

## Disclaimer

The research described in this article has been reviewed by the Center for Public Health and Environmental Assessment, U.S. Environmental Protection Agency, and approved for publication. Approval does not signify that the contents necessarily reflect the views and policies of the Agency, nor does the mention of trade names of commercial products constitute endorsement or recommendation for use. All opinions expressed in this paper are of the author’s and do not necessarily reflect the policies and views of DOE, or ORAU/ORISE.

## Acknowledgements

The authors thank Dr. M. Ian Gilmour of the U.S. EPA, Dr. Daniel L. Costa of the University of North Carolina (Formerly of the U.S. EPA) and Dr. Andrey Egorov of the U.S. EPA for their critical review of the manuscript. We acknowledge the help of Dr. Mark Higuchi and Mr. Abdul Malek Khan of the US EPA for ozone inhalation exposures.

## Author Contributions

A.R.H, S.J.S., U.P.K. designed experiments, data collection, and interpretation, manuscript preparation; T.W.J. data analysis, statistical evaluation, manuscript preparation; J.S.H., A.A.M.-R. data analysis and manuscript review; C.K.W.-C. data interpretation and manuscript review; M.C.S., D.I.A., C.N.M., A.K.F. M.S.H., R.G. data collection and manuscript review; A.J.G., D.D.-S., human study samples and manuscript review.

## REFERENCES

1. Landrigan PJ, Fuller R, Acosta NJR, et al. The Lancet Commission on pollution and health. Lancet. 2018;391(10119):462–512. doi:10.1016/S0140-6736(17)32345-0

2. Calderón-Garcidueñas L, Herrera-Soto A, Jury N, et al. Reduced repressive epigenetic marks, increased DNA damage and Alzheimer’s disease hallmarks in the brain of humans and mice exposed to particulate urban air pollution. Environ Res. 2020;183:109226. doi:10.1016/j.envres.2020.109226

3. Greve HJ, Mumaw CL, Messenger EJ, et al. Diesel exhaust impairs TREM2 to dysregulate neuroinflammation. J Neuroinflammation. 2020;17(1):351. doi:10.1186/s12974-020-02017-7

4. Paul KC, Jerrett M, Ritz B. Type 2 Diabetes Mellitus and Alzheimer’s Disease: Overlapping Biologic Mechanisms and Environmental Risk Factors. Curr Environ Health Rep. 2018;5(1):44–58. doi:10.1007/s40572-018-0176-1

5. Berman JD, Burkhardt J, Bayham J, Carter E, Wilson A. Acute Air Pollution Exposure and the Risk of Violent Behavior in the United States. Epidemiology. 2019;30(6):799–806. doi:10.1097/EDE.0000000000001085

6. Camargo Maluf F, Feder D, Alves de Siqueira Carvalho A. Analysis of the Relationship between Type II Diabetes Mellitus and Parkinson’s Disease: A Systematic Review. Parkinsons Dis. 2019;2019:4951379. doi:10.1155/2019/4951379

7. Norwitz NG, Mota AS, Norwitz SG, Clarke K. Multi-Loop Model of Alzheimer Disease: An Integrated Perspective on the Wnt/GSK3β, α-Synuclein, and Type 3 Diabetes Hypotheses. Front Aging Neurosci. 2019;11:184. doi:10.3389/fnagi.2019.00184

8. Hajat A, Diez Roux AV, Castro-Diehl C, et al. The Association between Long-Term Air Pollution and Urinary Catecholamines: Evidence from the Multi-Ethnic Study of Atherosclerosis. Environ Health Perspect. 2019;127(5):57007. doi:10.1289/EHP3286

9. Russell G, Lightman S. The human stress response. Nat Rev Endocrinol. 2019;15(9):525–534. doi:10.1038/s41574-019-0228-0

10. Herman JP, Nawreen N, Smail MA, Cotella EM. Brain mechanisms of HPA axis regulation: neurocircuitry and feedback in context Richard Kvetnansky lecture. Stress. 2020;23(6):617–632. doi:10.1080/10253890.2020.1859475

11. Biessels GJ, Despa F. Cognitive decline and dementia in diabetes mellitus: mechanisms and clinical implications. Nat Rev Endocrinol. 2018;14(10):591–604. doi:10.1038/s41574-018-0048-7

12. Danan D, Matar MA, Kaplan Z, Zohar J, Cohen H. Blunted basal corticosterone pulsatility predicts post-exposure susceptibility to PTSD phenotype in rats. Psychoneuroendocrinology. 2018;87:35–42. doi:10.1016/j.psyneuen.2017.09.023

13. Faulenbach M, Uthoff H, Schwegler K, Spinas GA, Schmid C, Wiesli P. Effect of psychological stress on glucose control in patients with Type 2 diabetes. Diabet Med. 2012;29(1):128–131. doi:10.1111/j.1464-5491.2011.03431.x

14. Teyin E, Derbent A, Balcioglu T, Cokmez B. The efficacy of caudal morphine or bupivacaine combined with general anesthesia on postoperative pain and neuroendocrine stress response in children. Paediatr Anaesth. 2006;16(3):290–296. doi:10.1111/j.1460-9592.2005.01711.x

15. Hásková A, Radovnická L, Petruželková L, et al. Real-time CGM Is Superior to Flash Glucose Monitoring for Glucose Control in Type 1 Diabetes: The CORRIDA Randomized Controlled Trial. Diabetes Care. 2020;43(11):2744–2750. doi:10.2337/dc20-0112

16. Baghelani M, Abbasi Z, Daneshmand M, Light PE. Non-invasive continuous-time glucose monitoring system using a chipless printable sensor based on split ring microwave resonators. Sci Rep. 2020;10(1):12980. doi:10.1038/s41598-020-69547-1

17. Torrente-Rodríguez RM, Tu J, Yang Y, et al. Investigation of cortisol dynamics in human sweat using a graphene-based wireless mHealth system. Matter. 2020;2(4):921–937. doi:10.1016/j.matt.2020.01.021

18. Miller DB, Karoly ED, Jones JC, et al. Inhaled ozone (O3)-induces changes in serum metabolomic and liver transcriptomic profiles in rats. Toxicol Appl Pharmacol. 2015;286(2):65–79. doi:10.1016/j.taap.2015.03.025

19. Miller DB, Ghio AJ, Karoly ED, et al. Ozone Exposure Increases Circulating Stress Hormones and Lipid Metabolites in Humans. Am J Respir Crit Care Med. 2016;193(12):1382–1391. doi:10.1164/rccm.201508-1599OC

20. Brockway R, Tiesma S, Bogie H, et al. Fully Implantable Arterial Blood Glucose Device for Metabolic Research Applications in Rats for Two Months. J Diabetes Sci Technol. 2015;9(4):771–781. doi:10.1177/1932296815586424

21. Henriquez AR, Snow SJ, Schladweiler MC, et al. Adrenergic and glucocorticoid receptor antagonists reduce ozone-induced lung injury and inflammation. Toxicol Appl Pharmacol. 2018;339:161–171. doi:10.1016/j.taap.2017.12.006

22. Miller DB, Snow SJ, Schladweiler MC, et al. Acute Ozone-Induced Pulmonary and Systemic Metabolic Effects Are Diminished in Adrenalectomized Rats. Toxicol Sci. 2016;150(2):312–322. doi:10.1093/toxsci/kfv331

23. Bass V, Gordon CJ, Jarema KA, et al. Ozone induces glucose intolerance and systemic metabolic effects in young and aged Brown Norway rats. Toxicol Appl Pharmacol. 2013;273(3):551–560. doi:10.1016/j.taap.2013.09.029

24. Henriquez AR, Williams W, Snow SJ, et al. The dynamicity of acute ozone-induced systemic leukocyte trafficking and adrenal-derived stress hormones. Toxicology. 2021;458:152823. doi:10.1016/j.tox.2021.152823

25. Henriquez AR, Snow SJ, Schladweiler MC, et al. Beta-2 Adrenergic and Glucocorticoid Receptor Agonists Modulate Ozone-Induced Pulmonary Protein Leakage and Inflammation in Healthy and Adrenalectomized Rats. Toxicol Sci. 2018;166(2):288–305. doi:10.1093/toxsci/kfy198

26. Griffin ÉW, Yssel JD, O’Neill E, et al. The β2-adrenoceptor agonist clenbuterol reduces the neuroinflammatory response, neutrophil infiltration and apoptosis following intra-striatal IL-1β administration to rats. Immunopharmacol Immunotoxicol. 2018;40(2):99–106. doi:10.1080/08923973.2017.1418882

27. Jonasson S, Wigenstam E, Koch B, Bucht A. Early treatment of chlorine-induced airway hyperresponsiveness and inflammation with corticosteroids. Toxicol Appl Pharmacol. 2013;271(2):168–174. doi:10.1016/j.taap.2013.04.037

28. Miller DB, Snow SJ, Henriquez A, et al. Systemic metabolic derangement, pulmonary effects, and insulin insufficiency following subchronic ozone exposure in rats. Toxicol Appl Pharmacol. 2016;306:47–57. doi:10.1016/j.taap.2016.06.027

29. Kainuma E, Watanabe M, Tomiyama-Miyaji C, et al. Association of glucocorticoid with stress-induced modulation of body temperature, blood glucose and innate immunity. Psychoneuroendocrinology. 2009;34(10):1459–1468. doi:10.1016/j.psyneuen.2009.04.021

30. Gordon CJ, Johnstone AF, Aydin C, et al. Episodic ozone exposure in adult and senescent Brown Norway rats: acute and delayed effect on heart rate, core temperature and motor activity. Inhal Toxicol. 2014;26(7):380–390. doi:10.3109/08958378.2014.905659

31. Hamade AK, Tankersley CG. Interstrain variation in cardiac and respiratory adaptation to repeated ozone and particulate matter exposures. Am J Physiol Regul Integr Comp Physiol. 2009;296(4):R1202–1215. doi:10.1152/ajpregu.90808.2008

32. Okada S, Yamaguchi N. Possible role of adrenoceptors in the hypothalamic paraventricular nucleus in corticotropin-releasing factor-induced sympatho-adrenomedullary outflow in rats. Auton Neurosci. 2017;203:74–80. doi:10.1016/j.autneu.2017.01.008

33. Henriquez AR, House JS, Snow SJ, et al. Ozone-induced dysregulation of neuroendocrine axes requires adrenal-derived stress hormones. Toxicol Sci. Published online August 9, 2019:kfz182. doi:10.1093/toxsci/kfz182

34. Lim CT, Khoo B. Normal Physiology of ACTH and GH Release in the Hypothalamus and Anterior Pituitary in Man. In: Feingold KR, Anawalt B, Boyce A, et al., eds. Endotext. MDText.com, Inc.; 2000. Accessed January 31, 2022. http://www.ncbi.nlm.nih.gov/books/NBK279116/

35. Qiu H, Yang JK, Chen C. Influence of insulin on growth hormone secretion, level and growth hormone signalling. Sheng Li Xue Bao. 2017;69(5):541–556.

36. Napolitano G, Barone D, Di Meo S, Venditti P. Adrenaline induces mitochondrial biogenesis in rat liver. J Bioenerg Biomembr. 2018;50(1):11–19. doi:10.1007/s10863-017-9736-6

37. Kodavanti UP. Susceptibility Variations in Air Pollution Health Effects: Incorporating Neuroendocrine Activation. Toxicol Pathol. 2019;47(8):962–975. doi:10.1177/0192623319878402

38. Snow SJ, McGee MA, Henriquez A, et al. Respiratory Effects and Systemic Stress Response Following Acute Acrolein Inhalation in Rats. Toxicol Sci. 2017;158(2):454–464. doi:10.1093/toxsci/kfx108

39. Gackière F, Saliba L, Baude A, Bosler O, Strube C. Ozone inhalation activates stress-responsive regions of the CNS. J Neurochem. 2011;117(6):961–972. doi:10.1111/j.1471-4159.2011.07267.x

40. Begg DP, Woods SC. Interactions between the central nervous system and pancreatic islet secretions: a historical perspective. Adv Physiol Educ. 2013;37(1):53–60. doi:10.1152/advan.00167.2012

41. Morgese MG, Schiavone S, Trabace L. Emerging role of amyloid beta in stress response: Implication for depression and diabetes. Eur J Pharmacol. 2017;817:22–29. doi:10.1016/j.ejphar.2017.08.031

42. Schatzberg AF, Lindley S. Glucocorticoid antagonists in neuropsychiatric [corrected] disorders. Eur J Pharmacol. 2008;583(2-3):358–364. doi:10.1016/j.ejphar.2008.01.001

43. Berger M, Sarnyai Z. “More than skin deep”: stress neurobiology and mental health consequences of racial discrimination. Stress. 2015;18(1):1–10. doi:10.3109/10253890.2014.989204

44. Snow SJ, Henriquez AR, Fisher A, et al. Peripheral metabolic effects of ozone exposure in healthy and diabetic rats on normal or high-cholesterol diet. Toxicol Appl Pharmacol. 2021;415:115427. doi:10.1016/j.taap.2021.115427

45. Goetsch VL, VanDorsten B, Pbert LA, Ullrich IH, Yeater RA. Acute effects of laboratory stress on blood glucose in noninsulin-dependent diabetes. Psychosom Med. 1993;55(6):492–496. doi:10.1097/00006842-199311000-00004

46. Stephens MAC, Mahon PB, McCaul ME, Wand GS. Hypothalamic-pituitary-adrenal axis response to acute psychosocial stress: Effects of biological sex and circulating sex hormones. Psychoneuroendocrinology. 2016;66:47–55. doi:10.1016/j.psyneuen.2015.12.021

47. Parra-Montes de Oca MA, Gutiérrez-Mariscal M, Salmerón-Jiménez MF, Jaimes-Hoy L, Charli JL, Joseph-Bravo P. Voluntary Exercise-Induced Activation of Thyroid Axis and Reduction of White Fat Depots Is Attenuated by Chronic Stress in a Sex Dimorphic Pattern in Adult Rats. Front Endocrinol (Lausanne). 2019;10:418. doi:10.3389/fendo.2019.00418

48. Kirby ED, Geraghty AC, Ubuka T, Bentley GE, Kaufer D. Stress increases putative gonadotropin inhibitory hormone and decreases luteinizing hormone in male rats. Proc Natl Acad Sci U S A. 2009;106(27):11324–11329. doi:10.1073/pnas.0901176106

49. Chatzitomaris A, Hoermann R, Midgley JE, et al. Thyroid Allostasis-Adaptive Responses of Thyrotropic Feedback Control to Conditions of Strain, Stress, and Developmental Programming. Front Endocrinol (Lausanne). 2017;8:163. doi:10.3389/fendo.2017.00163

50. Clemons GK, Wei D. Effect of short-term ozone exposure on exogenous thyroxine levels in thyroidectomized and hypophysectomized rats. Toxicol Appl Pharmacol. 1984;74(1):86–90. doi:10.1016/0041-008x(84)90273-4

51. Joseph-Bravo P, Jaimes-Hoy L, Uribe RM, Charli JL. 60 YEARS OF NEUROENDOCRINOLOGY: TRH, the first hypophysiotropic releasing hormone isolated: control of the pituitary-thyroid axis. J Endocrinol. 2015;226(2):T85–T100. doi:10.1530/JOE-15-0124

52. Colonna CH, Henriquez AR, House JS, et al. The Role of Hepatic Vagal Tone in Ozone-Induced Metabolic Dysfunction in the Liver. Toxicol Sci. 2021;181(2):229–245. doi:10.1093/toxsci/kfab025

53. Henriquez A, House J, Miller DB, et al. Adrenal-derived stress hormones modulate ozone-induced lung injury and inflammation. Toxicol Appl Pharmacol. 2017;329:249–258. doi:10.1016/j.taap.2017.06.009

54. Kuo T, McQueen A, Chen TC, Wang JC. Regulation of Glucose Homeostasis by Glucocorticoids. Adv Exp Med Biol. 2015;872:99–126. doi:10.1007/978-1-4939-2895-8_5

55. Patlar Akbulut F, Ikitimur B, Akan A. Wearable sensor-based evaluation of psychosocial stress in patients with metabolic syndrome. Artif Intell Med. 2020;104:101824. doi:10.1016/j.artmed.2020.101824

