## Supplemental Material 1 for "Glucose dynamics during ozone exposure measured using radiotelemetry: Stress drivers"

**Supplemental Material, Table S1.** A cross over experimental design for real time glucose monitoring used for animals implanted with telemeters.

| | | Rat | | | | | | |
|---|---|---|---|---|---|---|---|---|
| | | 1 | 2 | 3 | 4 | 5 | 7 | 8 |
| Week | 1st | 0.0 ppm | 0.0 ppm | 0.2 ppm | 0.2 ppm | 0.4 ppm | 0.8 ppm | 0.8 ppm |
| | 2nd | 0.8 ppm | 0.8 ppm | 0.0 ppm | 0.0 ppm | 0.2 ppm | 0.4 ppm | 0.4 ppm |
| | 3rd | 0.4 ppm | 0.4 ppm | 0.8 ppm | 0.8 ppm | 0.0 ppm | 0.2 ppm | 0.2 ppm |
| | 4th | 0.2 ppm | 0.2 ppm | 0.4 ppm | 0.4 ppm | 0.8 ppm | 0.0 ppm | 0.0 ppm |
| | 5th* | 0.8 ppm | 0.8 ppm | 0.8 ppm | 0.8 ppm | 0.0 ppm | 0.0 ppm | 0.0 ppm |
| | 6th* | 0.0 ppm | 0.0 ppm | 0.0 ppm | 0.0 ppm | 0.8 ppm | 0.8 ppm | 0.8 ppm |
| | 7th* | 0.8 ppm | 0.8 ppm | 0.8 ppm | 0.8 ppm | 0.0 ppm | 0.0 ppm | 0.0 ppm |

After a post-surgical wash-out period of 2 weeks concentration-dependent ozone effects were assessed. The actual number of exposures run over a period of four weeks for each concentration were n=6 for 0.0 ppm; n=5 for 0.2 ppm; n=5 for 0.4 ppm, and n=5 for 0.8 ppm. This is because of the issues with the probe collecting data for one rat. A cross-over design was used with one week washout period to allow assessment of different concentration related effects over a 4-week period. (1 rat had to be discarded [rat #6] due to problems in the surgical attachment of the glucose sensor. Note: On 05/30/2017, the ozone exposure was terminated at 3 hour 15 minutes due to ozone generation malfunction. *The 1-day exposure on 5th week began 4-days after the termination of 4th week exposure. Additional 1-day exposure began 5-days after week 5th exposure. Finally, week 7 exposure to air or 0.8 ppm ozone for 4 consecutive days began after 5 days of week 6 exposure.

**Supplemental Material, Table S2.** Demographic information of young healthy volunteers for a clinical study.

| Demographic | Data |
|---|---|
| Study participants | 34 |
| Sex | 16 males and 18 females |
| Age | 19-33 years |
| Race | 9-Blacks, 21-Whites, 3-Asians and 1-Hispanic |
| Sex Race ratio -White | 10 males and 11 females |
| Sex Race Ratio-Black | 3 males and 6 females |
| Sex Race Ratio-Asian | 3 males and 0 females |
| Sex Race Ratio-Hispanic | 0 male and 1 female |
| BMI (Mean ± SD)-overall | 25.2 ± 3.2 (n=34) |
| BMI (Mean ± SD)- Black female | 27.0 ± 3.4 (n=6) |
| BMI (Mean ± SD)- Black male | 27.4 ± 1.5 (n=3) |
| BMI (Mean ± SD)- White female | 23.9 ± 3.8 (n=11) |
| BMI (Mean ± SD)- White male | 24.6 ± 2.5 (n=10) |
| BMI (Mean ± SD)- Asian male | 27.2 ± 2.3 (n=3) |
| BMI (Mean ± SD)- Hispanic female | 23.3 ± 0.0 (n=1) |

Demographic information of volunteers participated in the clinical trial. No history of chronic disease, prescription medication use, and smocking.

**Supplemental Material, Figure S1.** Schematic of DSI glucose telemetry, placement of device in rat body and monitoring glucose levels in freely moving animals through signal receivers under each cage.

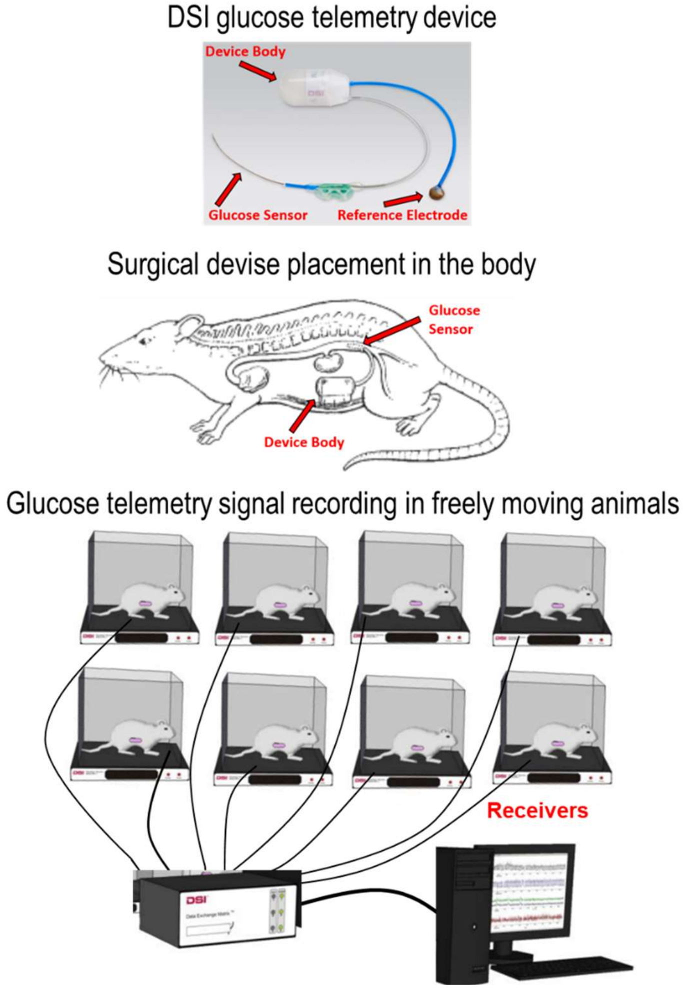


As explained in detailed methods above, rats were impanted with glucose telementry device and realtime glucose levels were monitored in individual home cages and in the air or ozone exposure chambers in freeely moving rats using DSI data processing and aquisition protocols (Brockway et al., 2015).

**Supplemental Material, Figure S2.** Ozone concentration-related changes in blood glucose and core body temperature during 4-hour exposure (0.0, 0.2, 0.4, and 0.8 ppm).

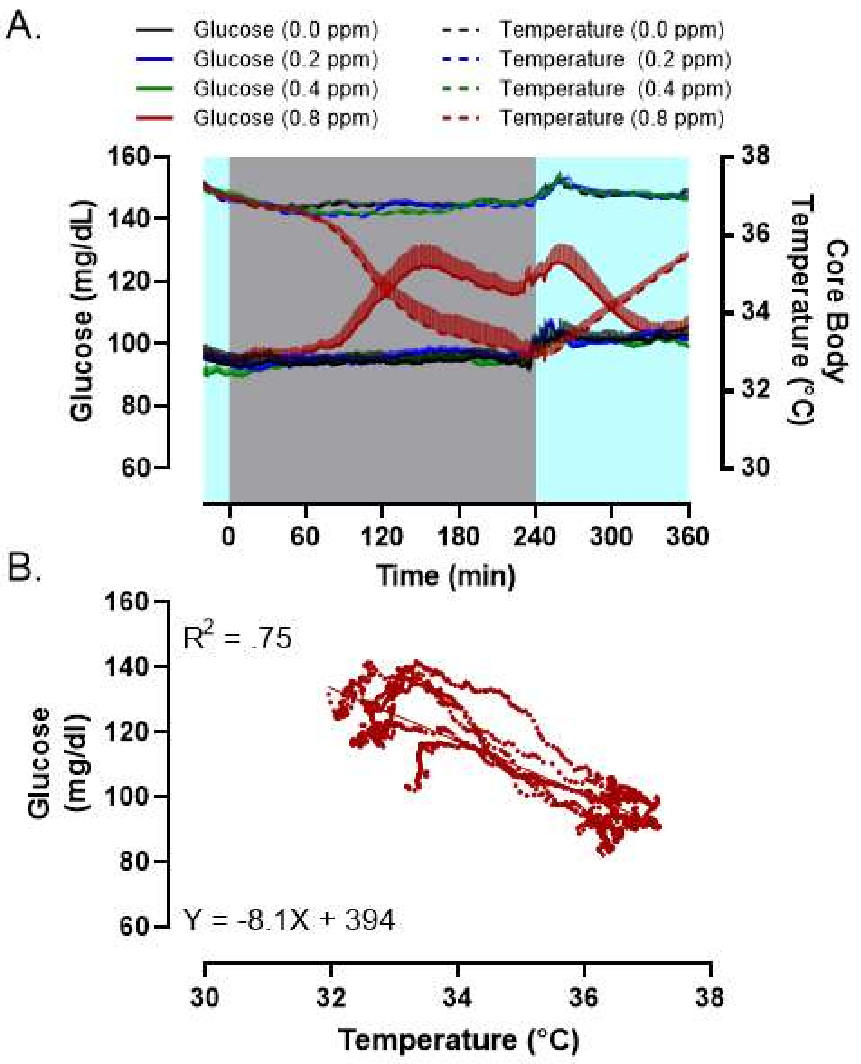


Using real-time glucose telemetry in rats, blood glucose levels and core body temperature were determined during and after ozone exposure. Glucose and temperature data averaged every minute are plotted as mean ± SEM of n=5-6, in a cross over design. Substantial temporal changes in circulating glucose and body temperature are noted only at 0.8 ppm ozone concentration. Temperature has a significant negative correlation with glucose noted only at 0.8 ppm ozone concentration.

**Supplemental Material, Figure S3**. Core body temperature changes during air or ozone exposures and non-exposure periods showing ozone effects and diurnal/nocturnal differences.

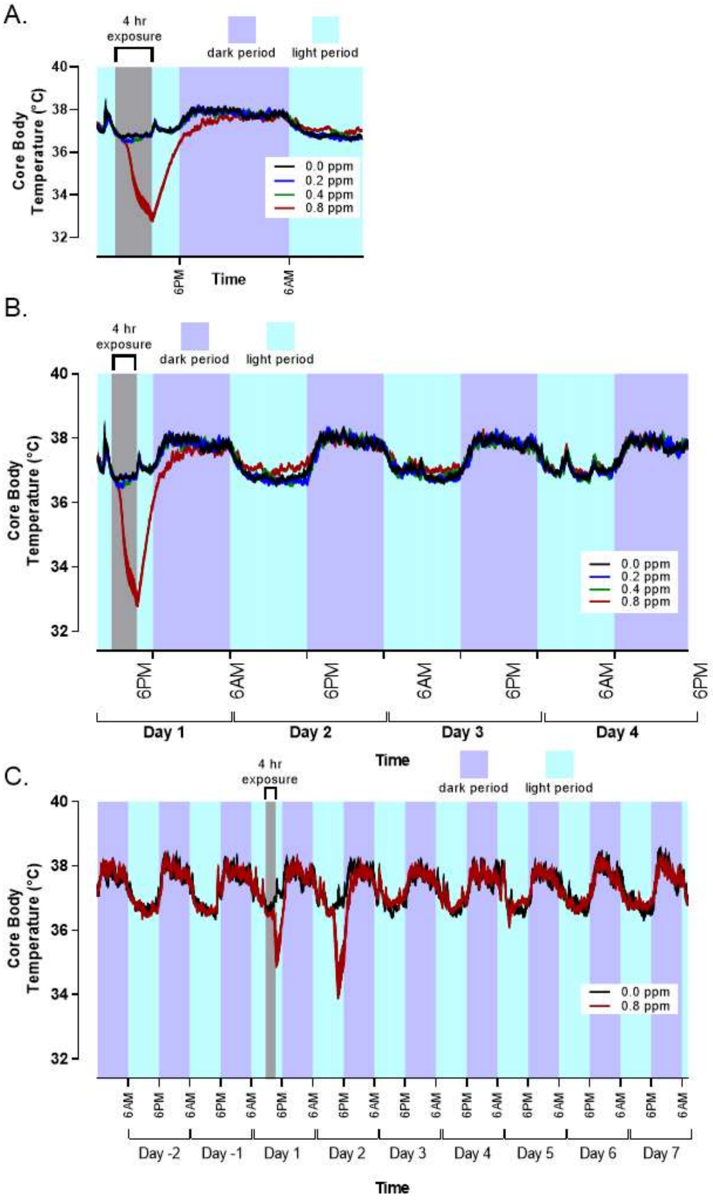


Using a cross over design, real-time core body temperature changes assessed by glucose telemetry device were computed during air and ozone exposure for 1-day (A), over 4-days (B) and 2-days prior to, during 4-consecutive days, as well as 3-days post-exposure recovery (C). Core body temperature data were averaged every minute and plotted as mean ± SEM of n=5-6.
